# Dynamic Filament Assembly Regulates the Prolyl Aminopeptidase Activity of Plant Immune Protein DM3

**DOI:** 10.64898/2026.09.03.749275

**Authors:** Nayun Kim, Wei-Lin Wan, Yi Yun Tan, In-Cheol Jang, Eunyoung Chae, Ji-Joon Song

## Abstract

The alpha/beta hydrolase DANGEROUS MIX 3 (DM3), a proline aminopeptidase in *Arabidopsis thaliana*, contributes to stress resilience through both metabolic regulation and immune signaling. Its functions are partitioned within its oligomeric structure as a trimer-of-dimers, in which the immune regulatory switch resides at the dimer interface. While the enzymatic activity of DM3 is not necessary for immune response, its activity to free prolines is critical to promote tolerance to salt and drought stress. However, how this activity is regulated has remained unclear. Here, we show that DM3 undergoes dynamic and reversible assembly into higher-order filaments in a salt-sensitive manner. The cryo-electron microscopy (EM) structure of filamentous DM3 reveals that the three-stranded helical filament arises from rearrangements of a planar hexamer into a tilted hexameric configuration. Notably, within these filaments, one of the catalytic residues is flipped out to disrupt the catalytic geometry, rendering the enzyme inactive. These findings establish filament assembly as a mechanism to sequester DM3 in an inactive state and suggest that the transition between distinct oligomeric states enables DM3 to coordinate its roles in biotic and abiotic stress responses.

**Significance statement:** Proteins with multiple interaction interfaces can assemble in different ways to regulate their function, but how this flexibility controls plant stress responses remains poorly understood. The *Arabidopsis* protein DM3 has dual roles as a metabolic enzyme and an immune regulator, and these functions are determined by its assembly state. Here, we show that DM3 reversibly assembles into higher-order filaments in a salt-sensitive manner. This transition disrupts its active site, switching off its enzymatic activity. Our findings reveal how structural plasticity in protein assemblies enables a single protein to coordinate multiple stress-response functions, highlighting a general mechanism for regulating protein activity in plants.

## Introduction

In plants, the diversification of immune receptors occasionally incurs the cost of autoimmunity, a phenomenon known as hybrid necrosis where mismatched immune components trigger inappropriate defense responses (1, 2). DANGEROUS MIX 3 (DM3), an alpha/beta hydrolase, triggers an immune response together with the nucleotide-binding leucine-rich repeat (NLR) protein DM2 that is encoded by the most rapidly evolving gene cluster in *Arabidopsis thaliana* (hereafter, *Arabidopsis*) (3, 4). We previously determined cryo-EM structures of an autoimmune-triggering DM3 variant (DM3^Hh-0^) and its non-triggering counterpart (DM3^Col-0^), revealing that both predominantly assemble into hexameric complexes with a trimer-of-dimers configuration in D3 symmetry (5). In that study, we demonstrated that the DM3 allele from the *Arabidopsis* accession Hh-0 functionally interacts with DM2h from Bla-1 to induce immunity, and that the structural integrity at the DM3 dimer interface is critical for this immune activation while the catalytic activity is not required.

Distinct from its role as an immune component, DM3 intrinsically functions as a prolyl aminopeptidase 2 (PAP2), playing a critical role in proline metabolism by cleaving the N-terminal proline residue from a peptide to replenish the cellular proline pool in mitochondria (6). In plants, proline functions not only as a fundamental building block for proteins but also as a multifunctional metabolite important for environmental adaptation (7, 8). Under abiotic stress conditions, such as high salinity and drought, the accumulation of proline is a primary defense mechanism (6, 9–11). Proline functions as a compatible osmolyte, which accumulates to high levels without disrupting proteins or metabolism, to maintain cell turgor, serves as a potent antioxidant to scavenge reactive oxygen species (ROS) and stabilize protein structures (7, 8, 10, 12). As a result, disruption of proline metabolism leaves plants highly vulnerable to stress, compromising their growth and survival (13–15). Therefore, fine-tuning proline homeostasis through biosynthetic, catabolic, and recycling pathways is critical for stress tolerance. Previous studies using *DM3/PAP2* knockout *Arabidopsis* mutants have demonstrated that loss of the catalytic function of DM3 results in reduced proline content and increased susceptibility to salt and drought stress (6).

Remarkably, the highly organized structural assembly of DM3 serves as the critical factor compartmentalizing its dual functions in immunity and metabolism. Specifically, the hexamer is assembled via two distinct interaction interfaces, the dimer and trimer interfaces, which establish a division of labor within the complex being required for immune activation and metabolic regulation, respectively. The structural modularity of these distinct interfaces highlights the dynamic nature of DM3. Furthermore, our previous *in planta* BN-PAGE analyses revealed a slow-migrating band above the expected hexameric complex (5). This observation raised the question of whether additional, higher-order oligomeric states exist to regulate this protein *in vivo*.

At the supramolecular level, filament formation has emerged as a regulatory strategy that enables metabolic enzymes and immune receptors to modulate their activity in response to biochemical and physicochemical conditions (16–19). Filaments can suppress activity by constraining active conformations or sequestering functional domains. Bacterial CTP synthase (CTPS), yeast glucokinase 1 (Glk1), glutamine synthetase 1 (Gln1), and the plant immune receptor *Solanum lycopersicum* NLR required for cell death 2 (SlNRC2) represent such inhibitory assemblies (20–23). Conversely, filaments can enhance activity. For instance, human acetyl-CoA carboxylase (ACC) and CTPS stabilize active states upon filamentation, and bacterial aldehyde-alcohol dehydrogenase (AdhE) and delta-1-pyrroline-5-carboxylate synthase from *Arabidopsis* (AtP5CS) form internal channels for efficient substrate transfer (24–27). Moreover, some proteins are repurposed upon filamentation, acquiring new functions. The Toll/interleukin-1 receptor (TIR) domain of the L7 plant immune receptor and 2-Cys peroxiredoxin (Prx) transition from an NADase and an antioxidant enzyme to a 2’,3’-cAMP/cGMP synthetase and a molecular chaperone, respectively (28, 29).

To determine whether DM3 utilizes a comparable supramolecular strategy, we aimed to characterize the assembly dynamics of DM3 related to its function. Here, we show that DM3 undergoes a dynamically reversible transition into inhibitory filaments upon trimer interface disassembly, serving as a negative regulatory mechanism to silence its enzymatic activity.

## Results

### DM3 forms into filamentous structure

We previously showed that non-autoimmune DM3 variant from the *Arabidopsis* Col-0 accession (residues 52-515, DM3_WT_) adopts a planar hexameric trimer-of-dimers configuration, with dimeric and trimeric interfaces contributing to different functions (5). To assess the contribution of the trimer interface to DM3 assembly, we generated a trimer destabilizing mutant (DM3_DT_) by truncating the last five C-terminal amino acids that are inserted into the pocket at the trimer-interface within the hexamer (5). Recombinant DM3_DT_ expressed in *Escherichia coli* (*E. coli*) exhibited reduced trimer-interface stability and eluted at a dimeric position in size-exclusion chromatography (SEC) (*SI Appendix*, Fig. S1*A*).

However, unexpectedly, negative-stain electron microscopy (EM) analysis showed filamentous DM3 in addition to the dimeric DM3, indicating that DM3_DT_ assembled into filamentous structures (Fig. 1*A*). The filaments likely formed from the dimeric population of DM3_DT_ during the interval between SEC and negative-stain EM sample preparation. To confirm that filament formation is induced by compromise of the trimer interface, rather than being an artifact of the C-terminal deletion, we generated another mutant, DM3 Y460E/V484E (DM3_YV-EE_), designed to destabilize the trimer interface based on our previous structure of DM3 hexamer (5). We observed that the DM3_YV-EE_ also forms filaments (Fig. 1*B* and *SI Appendix,* Fig. S1*B*), albeit to a much lesser extent in frequency and length. Taken together, our observations demonstrate that the destabilization of the trimer induces the filament formation.

**Fig. 1.**
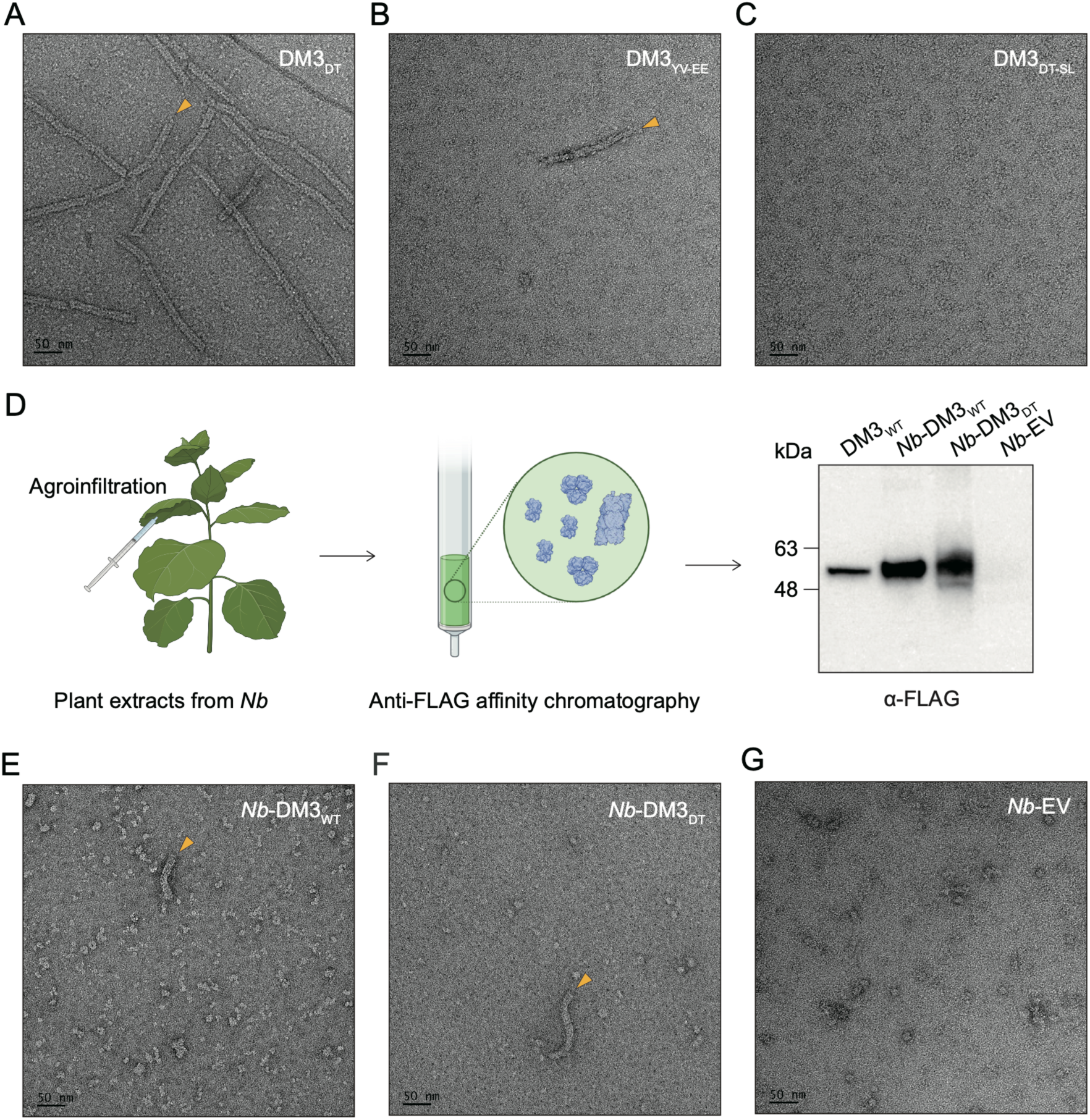
Filament formation of DM3. *(A, B, and C)* Representative negative-stain EM images of DM3_DT_ *(A)*, DM3_YV-EE_ *(B)*, and DM3_DT-SL_ *(C)*. *(D)* A schematic illustration of the transient expression of DM3 variants in *Nicotiana benthamiana* (*Nb*) leaves via agroinfiltration, followed by the purification of FLAG-tagged DM3 variants using anti-FLAG affinity chromatography. The purified samples were analyzed by western blot. Recombinant DM3_WT_ purified form *E. coli* was used as a standard. Samples include *Nb*-expressed DM3_WT_ (*Nb*-DM3_WT_) and DM3_DT_ (*Nb*-DM3_DT_), with an empty vector (EV) serving as a negative control. *(E, F, and G)* Representative negative-stain EM images of *Nb*-DM3_WT_ *(E)*, *Nb*-DM3_DT_ *(F)* and *Nb*-EV *(G)*. Filamentous structures are highlighted with yellow arrowheads in all panels.

We next examined whether plant-expressed DM3 also forms filaments. We transiently expressed FLAG-tagged wild-type DM3 (DM3_WT_) and DM3_DT_ under the 35S promoter in *Nicotiana benthamiana* (*Nb*) leaves. The expression and purification of these proteins via anti-FLAG affinity chromatography were confirmed by western blot (Fig. 1*D*). Negative-stain EM analysis of DM3 expressed in *Nb* showed the presence of filaments in both DM3_WT_ (Fig. 1*E*) and DM3_DT_ (Fig. 1*F*) samples, which existed as a minor population alongside various other oligomeric configurations, while no filaments were observed in the sample infiltrated with an empty vector (EV) (Fig. 1*G*). It is notable that the plant-expressed DM3_WT_ forms filaments, while the filament formation of recombinant DM3_WT_ was not apparent (5). This can be attributed to the processing of DM3_WT_ molecules in the plant cell (5), which potentially generates a fraction of DM3 molecules experiencing the destabilization of the trimer interface, such as C-terminal cleavage or else, favoring filament formation.

### The cryo-EM structure of filamentous DM3

To elucidate the detailed molecular architecture of the DM3 filament, we determined the cryo-EM structure of DM3_DT_ at 3.3 Å resolution by helical processing (Fig. 2 and *SI Appendix*, Fig. S2 and S3). We vitrified filamentous DM3_DT_ and collected 14,658 micrographs using a Titan Krios 300 keV microscope. The 2D class averaging of the particles revealed that DM3_DT_ exhibited a heterogeneous population from dimers, hexamers and filaments (Fig. 2*A* and 2*B*). To obtain the atomic resolution structure of the filamentous DM3_DT_, we extracted particles of a filamentous form and further processed to obtain 3D volume of the filamentous DM3_DT_ (Fig. 2*D*). The atomic resolution cryo-EM structure delineated that the filament adopts a triple helical architecture. Within this assembly, one helical turn pitch per single strand is composed of 16 asymmetric units along a left-handed screw axis (Fig. 2*C*). Each asymmetric unit corresponds to a dimeric DM3, with a twist of −22.5° and a rise of 63.9 Å calculated from the helical processing. In the filament, dimeric DM3 are radially arranged around the helical axis, forming a hollow assembly with an outer diameter of approximately 140 Å and an inner hole diameter of 35 Å (Fig. 2*C* and 2*E*). As a result of the 3-start helical packing, three dimers forming a tilted hexameric form are present within a single cross-sectional plane. We compared the six-subunit arrangement of filamentous DM3 and the previously reported planar hexameric DM3 (PDB: 8JGN) (5). This comparison revealed that a tilted hexameric assembly forms the basis of the filament with a −22.5° helical twist away from the planar hexameric configuration that DM3_WT_ adopts (Fig. 2*F* and 2*G*). The remarkably low RMSD of 0.729 Å between the monomer units extracted from the filament and hexamer suggests the structural rearrangement from the planar hexamer to the filament is not primarily driven by major conformational changes within individual monomers (*SI Appendix*, Fig. S4*A*), but rather by a reconfiguration of the intermolecular interactions among DM3 subunits. Consistent with this, the RMSD values between two adjacent DM3 subunits at the dimer interface and at the trimer interface were 1.194 Å (*SI Appendix*, Fig. S4*B*) and 5.922 Å (*SI Appendix*, Fig. S4*C*), respectively. The low RMSD at the dimer interface indicates that the dimeric unit is structurally preserved, whereas the high RMSD at the trimer interface reflects that a reorientation occurs between the dimeric building blocks (at the trimer interface) during filament assembly. Taken together with our previous work on the planar hexameric structure (5), this cryo-EM structure shows that DM3 undergoes a dynamic structural transition in its higher-order assembly.

**Fig. 2.**
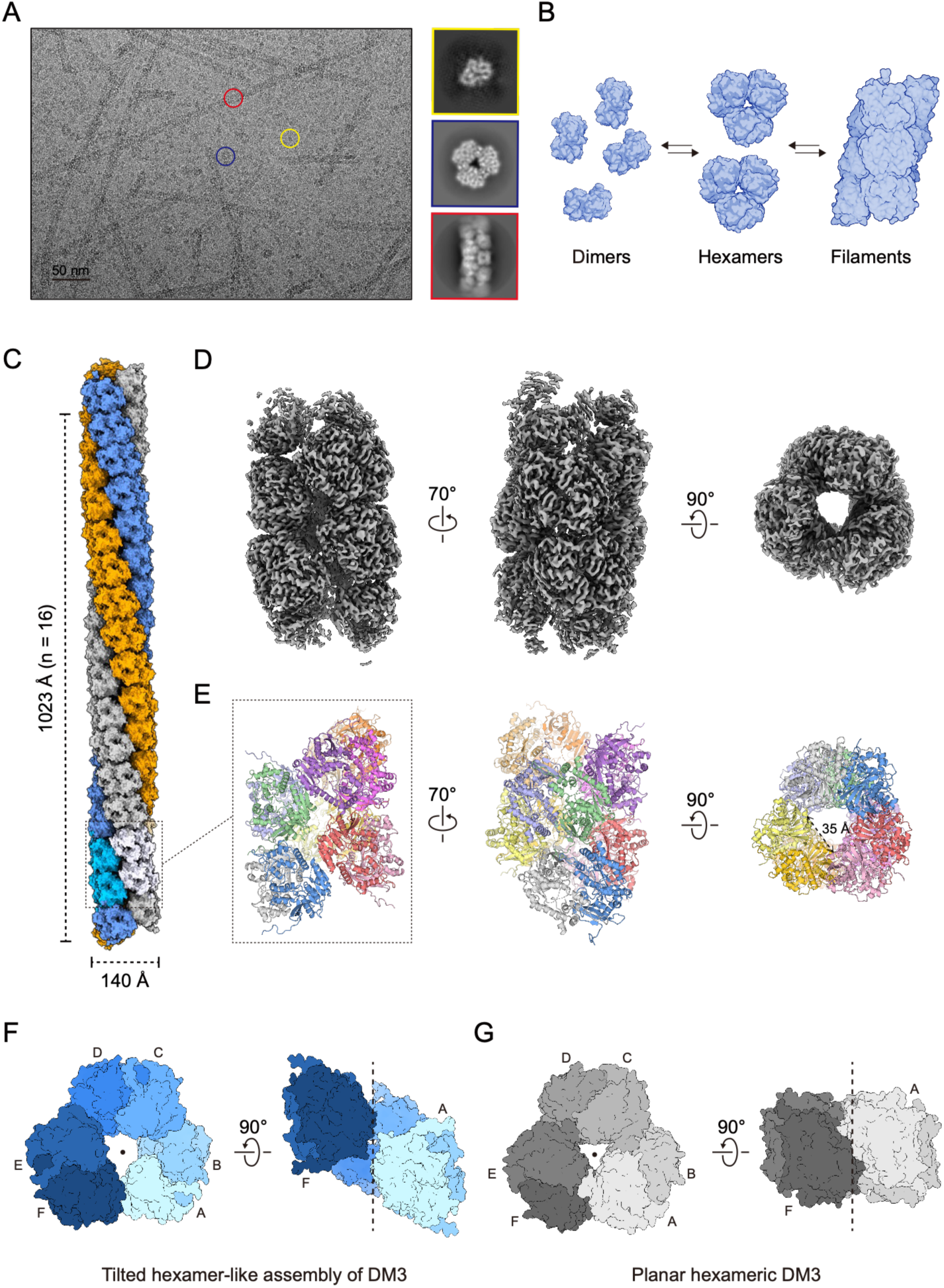
Cryo-EM structure of filamentous DM3_DT_. *(A)* Cryo-EM micrograph shows a heterogeneous population of DM3_DT_ oligomers including dimers (yellow), hexamers (blue), and filaments (red). *(B)* Schematic model of the structural equilibrium of DM3_DT_. *(C)* Surface representation shows one helical turn of DM3_DT_ filament. Individual protofilaments are differentially colored. The dotted box highlights the specific segment used for the cryo-EM data processing, corresponding to two tilted hexameric layers. *(D)* The cryo-EM map of DM3_DT_ filament. *(E)* The atomic resolution cryo-EM structure of DM3_DT_ filament represented as a cartoon, showing the side, front, and top views from left to right. Individual chains are differentially colored. *(F)* Top (left) and side (right) views of a tilted hexameric unit in a DM3_DT_ filament. The filament axis is indicated by a black dot in the top view and a dashed line in the side view. *(G)* Top (left) and side (right) views of a six-subunit in a planar DM3_WT_ hexamer.

### Reconfiguration of the trimer interface and the N-terminal loop in DM3 filament assembly

The atomic resolution structure allowed us to examine the detailed interactions critical for filament formation (Fig. 3*A* and 3*B*). The dimer interfaces of DM3_DT_ are highly similar to the interfaces observed in the planar hexameric structure (Fig. 3*C* and *SI Appendix*, Fig. S4*D* and S4*F*) (5). The dimer interface is mediated by interactions between the dimerizing loop (DL; residues 329-344) and the dimerizing helix (DH; residues 166-171), which we previously defined in the planar hexameric structure (Fig. 3*C*) (5). Dimers further associate laterally with adjacent dimers through hydrophobic interactions on the C-terminal region of the ABH domain at the trimer interface (Fig. 3*D*). At this interface, the β8 strands are arranged in an antiparallel manner, bringing hydrophobic residues, including V484 on β8, Y460 on the loop connecting β7 and α5, and F467 on α5, into close proximity. Compared with the trimer interface in the planar hexameric structure (*SI Appendix*, Fig. S4*G*), the hydrophobic interactions are weakened in the filament, with increased distances between the residues. This reconfiguration at the trimer interface renders DM3 into a tilted open-ring configuration. The subsequent stacking between the tilted hexameric assemblies is mediated by the N-terminal loop (residues 56-68), hereafter referred to as the Stacking Loop (SL) (Fig. 3*A* and 3*E*). E64 of the SL forms a salt bridge with K66 of the SL from the axially adjacent layer, therefore forming double inter-loop salt bridges between the SLs that mediate stacking assembly (Fig. 3*E* and *SI Appendix*, Fig. S4*E*). In the planar hexameric assembly, the cryo-EM map corresponding to the SL is absent, indicating that this region becomes ordered in the filamentous structure (5). In the filamentous structure, a deep pocket formed by the DH and DL accommodates the SL, primarily via back-bone interactions (Figures 3*F*, 3*G* and *SI Appendix*, Fig. S5). Specifically, the main chain of S56 of the SL forms a hydrogen bond with E177, and the main chain of D60 and G63 of the SL form hydrogen bonds with the main chain of K99 and the hydroxyl group of T101, respectively. Additionally, Y58 of the SL is positioned in close proximity to the hydrophobic cluster previously described at the dimer interface (Fig. 3*F*). To investigate the contribution of the SL for filament formation, we deleted the SL (DM3_DT-SL_) and examined its oligomeric state by size-exclusion chromatography followed by negative-stain EM (Fig 1*C* and *SI Appendix*, Fig. S1*C*).

**Fig. 3.**
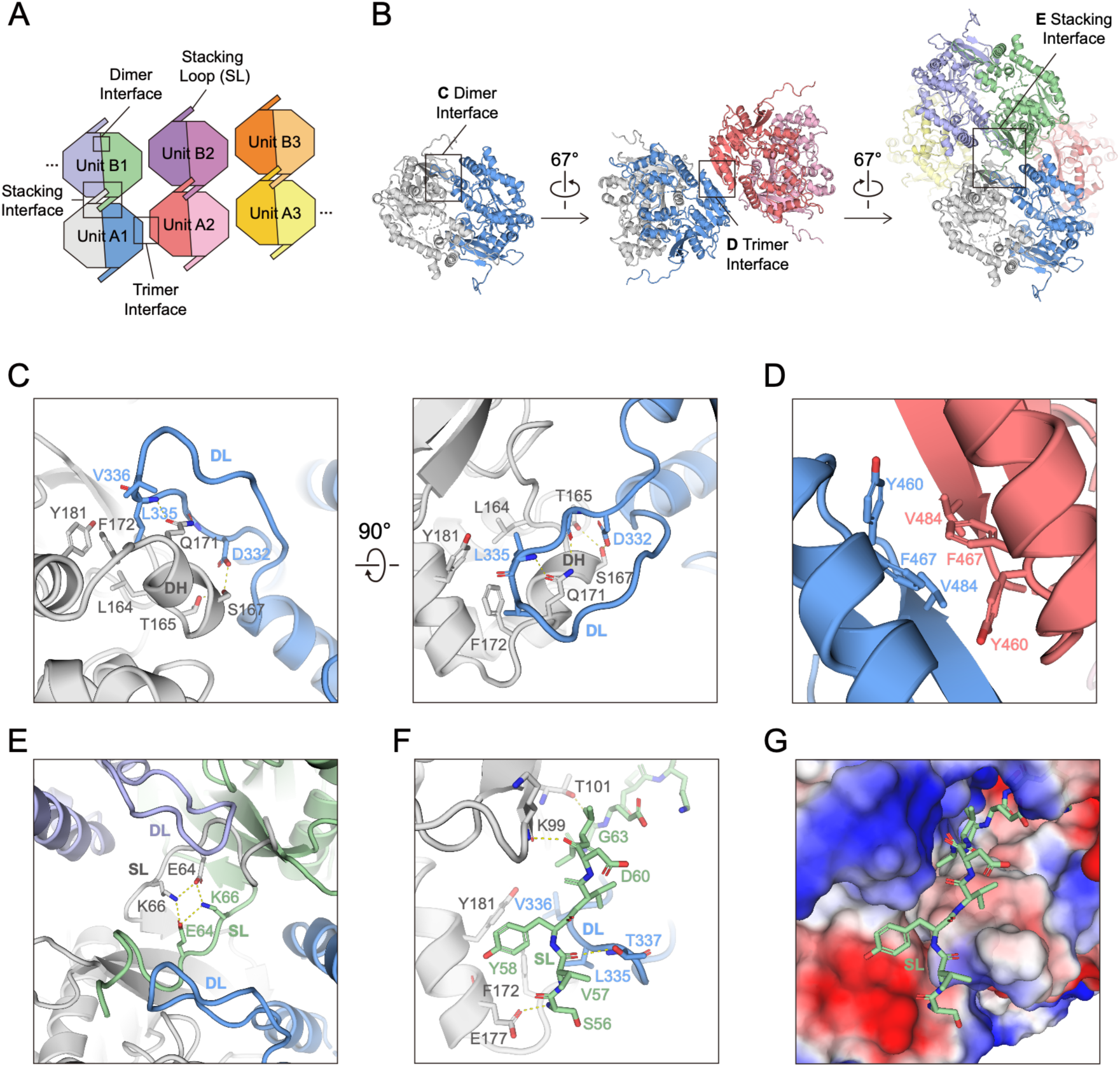
Structural determinants of the DM3_DT_ filament formation. *(A)* Schematic representation of filament architecture highlighting dimer, trimer, stacking interfaces and stacking loop, depicted in a linearized 1D arrangement. The dimeric units are grouped alphabetically to clearly illustrate the tilted hexameric configuration, with Units A1, A2, and A3 forming the first tilted hexameric layer, and Units B1, B2, and B3 constituting the subsequent layer. The interaction between Unit A3 and Unit B1 through trimer interface is depicted by the dotted line. *(B)* Dimer, trimer, and stacking interface are shown in the cryo-EM structure of DM3_DT_ filament. Each interface is indicated by a box. *(C)* Detailed view of dimer interface formed by DH (residues 166-171) and DL (residues 329-344). *(D)* Detailed view of trimer interface formed by subunits arranged in an antiparallel manner, with close hydrophobic interactions. *(E)* Detailed view of stacking interface. The inter-loop interactions are stabilized by salt bridges between K66 and E64. *(F)* Insertion of the SL into the axially adjacent dimer interface through backbone interactions and the interaction of Y58 with a hydrophobic cluster at the dimer interface are shown. *(G)* The SL is shown as a stick, and the dimer interface is displayed as a surface colored according to electrostatic potential (−50 to +50 mV), with red indicating negative and blue indicating positive potential.

Consistent with these structural observations, the deletion of this loop (DM3 68-510) abolishes filament assembly. Our cryo-EM structure reveals the structural versatility of DM3, which can adopt various oligomeric states ranging from dimers to filaments.

### Filament formation of DM3 rearranges the catalytic triad, leading to the loss of the activity

DM3 is a proline aminopeptidase with a conserved catalytic triad composed of S214, H490 and D462 (5). We previously showed that the catalytic triad is compartmentalized inside of the planar hexameric structure and the structural integrity is required for the PAP activity. Interestingly, our structural analysis revealed that the filamentation of DM3 reorganize the conformation of the catalytic triad. While S214 remains compartmentalized within the barrel and located at the same position to that of the planar hexamer (Fig. 4*A*), D462 flips out toward and is located at the outer surface of DM3 (Fig. 4*B*). In addition, the region containing H490 is highly disordered. This observation indicates that the filament formation of DM3 leads to an inactive form. These data suggest that the oligomeric states of DM3 affect the enzymatic activity. Therefore, we compared the enzymatic activities of DM3s having different oligomeric equilibrium state. We measured the PAP activity of DM3_WT_, DM3_DT_, and DM3_DT-SL_ (Fig. 4*C*). DM3_WT_, which forms a stable planar hexameric structure, showed robust PAP activity, consistent with our previous study (5). DM3_DT_, which exists as dimers and hexamers but predominantly adopts a filamentous form, as shown cryo-EM analysis (Fig. 2*A* and 2*B*), exhibited greatly reduced activity, with residual activity remaining. This data is consistent with the structure of the filamentous DM3 where the catalytic triad is disrupted.

**Fig. 4.**
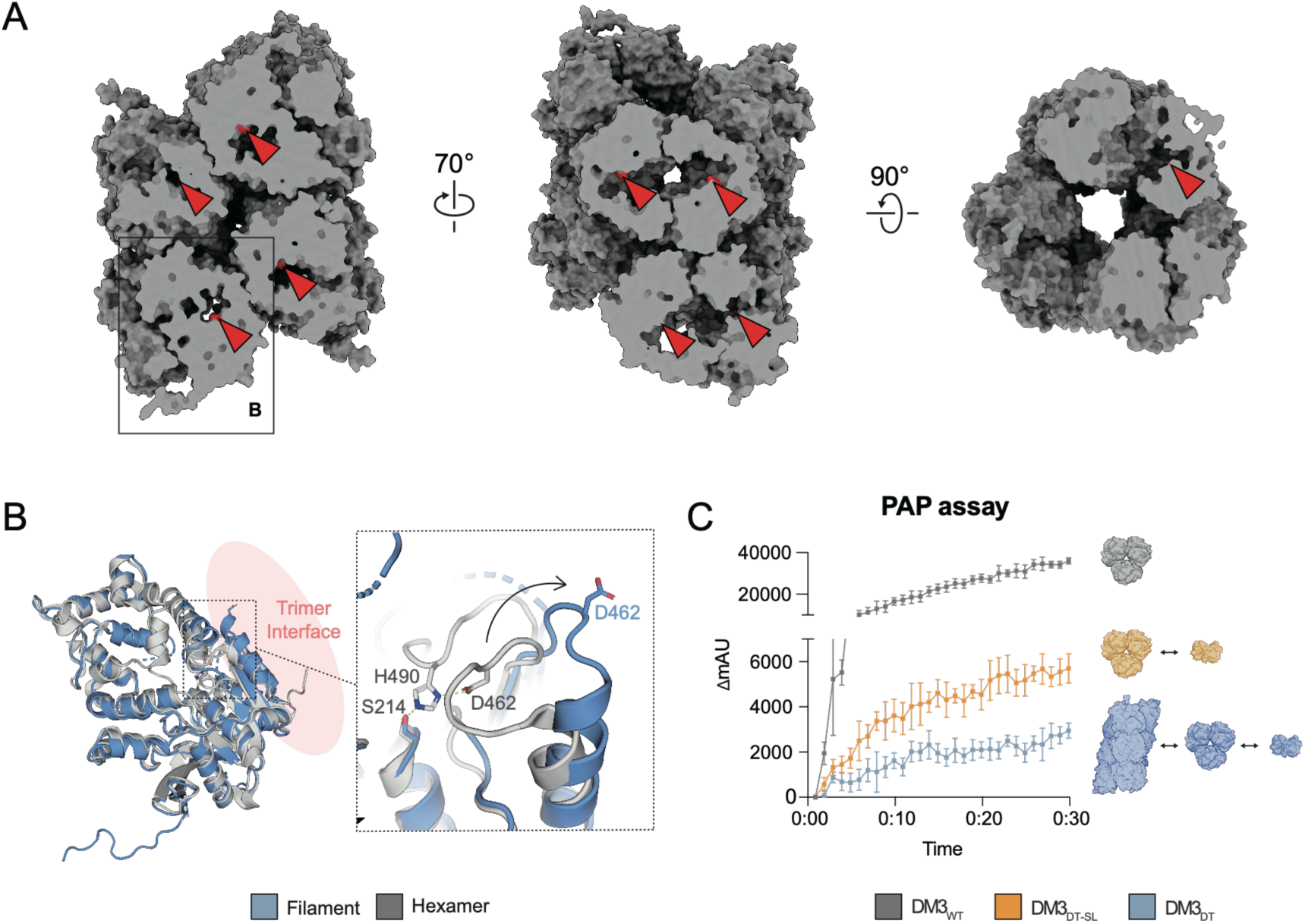
Disruption of the catalytic triad in the DM3_DT_ filament. *(A)* Locations of S214 residues in DM3_DT_ filament. Catalytic residue S214 is colored red. Red arrowheads point to catalytic triad residues (S214) within the filament structure in each orientation. *(B)* Single chains from the hexamer (gray) and filament (blue) were superimposed. A detailed view of catalytic triad is shown. In filament, D462 is flipped out toward trimer interface, as indicated by the arrow. *(C)* PAP activity assay of DM3 variants.

The residual activity is likely from the existing hexameric form of DM3_DT_. DM3_DT-SL_ can form dimers and hexamers, but does not form filaments due to the absence of the SL. This filament-deficient state resulted in two-fold higher activity than DM3_DT_, although its activity remained much lower than that of DM3_WT_. These data further support that filamentous form of DM3 represent a strongly disabled configuration regarding to the catalytic activity. Our *in vitro* PAP activity assay together with the structural analysis suggests that the oligomeric states govern the enzymatic activity of DM3.

### Osmotic condition alters the DM3 filament formation

As PAP activity is crucial to handle osmotic stress such as high salinity (6) and our data showed that the filamentous DM3 is an inactive form, we investigated whether the ionic strength affects the degree of filament formation of DM3. We found that DM3_DT_ stay as a non-filamentous form at 0 mM salt (*SI Appendix*, Fig. S6). We incubated the non-filamentous DM3_DT_ in the presence of several solutes including NaCl, KCl and mannitol at 50, 100, 200, 400 mM concentrations and examined the filament formation (Fig. 5*A* and 5*B*). We found that filament assembly was most extensive at 50 mM NaCl or KCl, and the filament assembly decreased progressively as the salt concentration increased. The filament assembly was completely abolished at 200 mM and above concentration of both salts. In contrast, DM3 does not form filamentous structure in the presence of mannitol under the condition used.

**Fig. 5.**
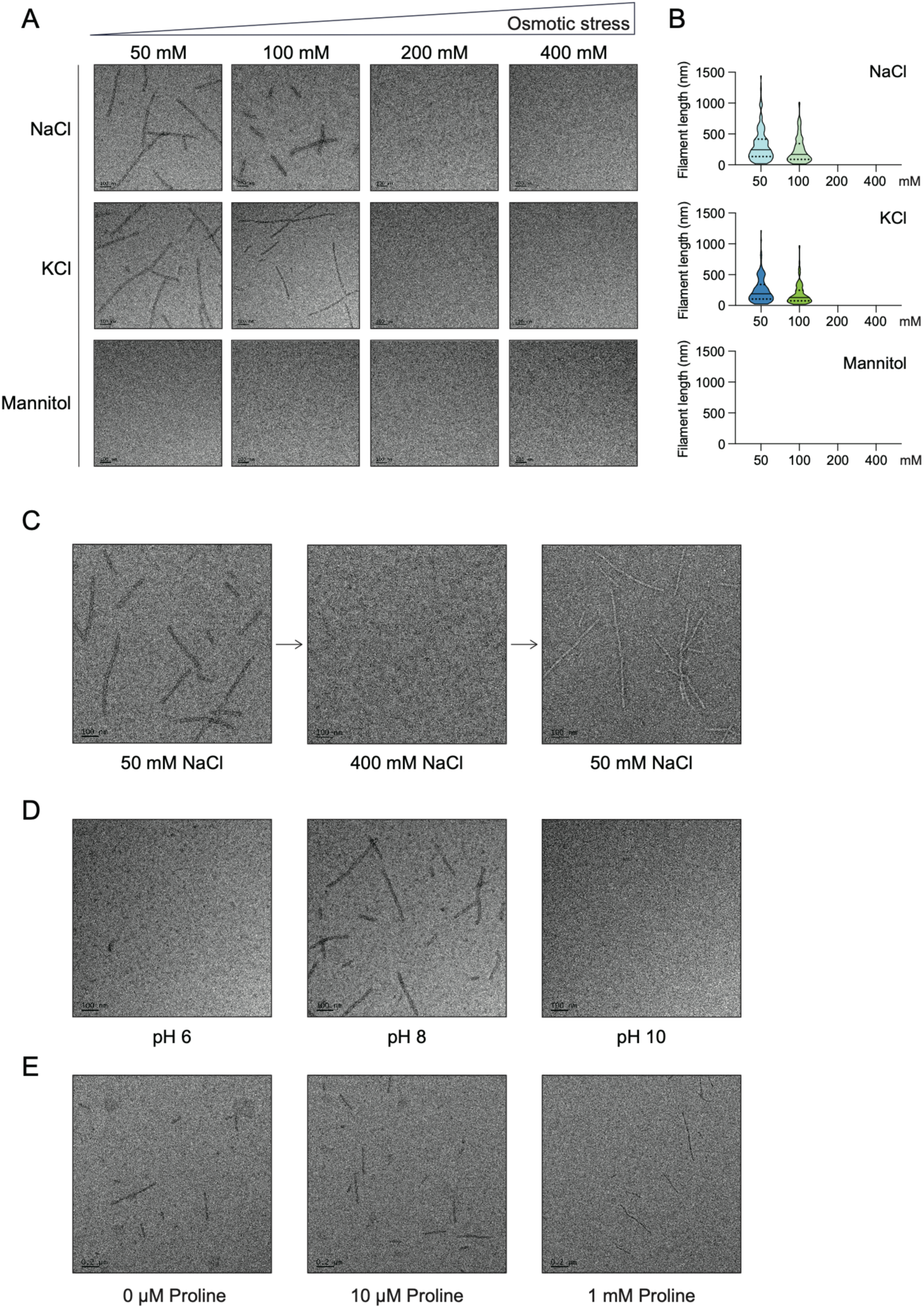
Dynamics of DM3 filament formation. *(A)* Effect of osmotic stress on filament assembly. Negative-stain EM images show DM3_DT_ filament states increasing concentrations (50-400 mM) of NaCl, KCl, and mannitol. *(B)* Violin plots show the distribution of filament states at varying concentration of NaCl, KCl, and mannitol. Solid and dashed lines represent medians and quartiles, respectively. *(C)* Salt-dependent reversibility of DM3_DT_ filament assembly. Filaments assembled at 50 mM NaCl disassembled at 400 mM NaCl and reassembled upon returning to 50 mM NaCl. *(D)* Representative negative-stain EM images of DM3_DT_ at pH 6, 8, and 10 are shown. *(E)* Representative negative stain EM images of DM3_DT_ are shown for the negative control (left), and after addition of 10 µM (middle) and 1 mM (right) proline.

These data suggest that filament assembly of DM3 is specifically responsive to ionic strength rather than general osmotic stress. We next tested whether the DM3 filamentation can be reversed by changing the salt concentration (Fig. 5*C*). The DM3 filaments formed at 50 mM NaCl disappeared after buffer exchange to 400 mM NaCl. Upon returning to 50 mM NaCl, the DM3 filaments reformed, demonstrating the reversibility of DM3 filamentation. We also investigated the effect of pH on filament formation. We found that filament assembly occurs only at near-physiological pH (pH 8.0) (Fig. 5*D*). Furthermore, as DM3 is a proline aminopeptidase that generates free-proline as a product, we tested whether the product itself modulates the assembly process. The addition of proline resulted in no significant difference in filament formation, indicating that assembly is not regulated by its enzymatic product (Fig. 5*E*). These data suggest that the filament formation is dynamic process, which can be regulated by environmental conditions.

## Discussion

Our findings establish higher-order assembly as a regulatory principle controlling DM3 activity. Together with our previous work (5), we show that DM3 exists in a dynamic equilibrium spanning monomeric, dimeric, hexameric and filamentous states. The dimer serves as the fundamental building block, assembling laterally into a finite, planar trimer-of-dimers hexamer or engaging in iterative multivalent interactions to generate a three-stranded filament through coordinated lateral and axial contacts. Disruption or weakening of the trimer interface within the structurally defined hexamer drives the transition toward filament formation, converting a closed oligomer into an open-ended multivalent polymer. This reconfiguration repurposes pre-existing interfaces to make them available for stacking loop (SL) binding, and the SL then inserts into the adjacent dimer interface surrounding the DH– DL pocket and nucleates axial assembly. Filament formation is directly coupled to catalytic silencing. Repositioning of two catalytic triad residues, Asp (D) and His (H), away from the nucleophilic serine disrupts active-site geometry, while SL insertion further occludes the DH– DL pocket. Because structural integrity of this DH-DL pocket functions as a previously defined immune switch (5), filament assembly may not only stabilize an autoinhibited catalytic configuration but also may reinforce immune restraint. In this way, the filamentous architecture demonstrates how multivalent supramolecular organization governs both enzymatic competence and immune potentiation within a single bifunctional enzyme complex. However, at this moment, it is not clear whether the filament assembly is directly related with plant immunity. Further study will be required to understand the role of filament assembly, if any, in regulating immune response.

The DM3 assembly switch is reversible and responsive to ionic conditions. Filament stability depends on the reconfigured trimer and stacking interfaces, and the multivalent polymer’s sensitivity to ionic strength is likely governed by the electrostatic interactions at the stacking interface. Elevated salt conditions favour disassembly, which would shift the equilibrium toward the active hexamer, whereas restoration of homeostasis promotes filament formation and enzymatic silencing. Rather than functioning as a static autoinhibitory sink, the filament therefore may represent an ion-sensitive structural rheostat, raising a possibility that DM3 assembly could be dynamically tuned by changes in ionic conditions. This reversible assembly behavior aligns with DM3’s role in salinity tolerance through regulation of proline, a key osmolyte in plants (6). Under high-salt stress, a restoration of the hexameric state from the autoinhibitory filament would sustain proline levels on demand, whereas subsequent filament formation could prevent excessive osmolyte accumulation once stress subsides.

Similar environmentally responsive assembly mechanisms have been observed in other regulatory proteins (e.g., casein kinase 2, caspase-9 CARD) (30, 31), supporting the broader principle that higher-order organization provides a strategy for metabolic control. Notably, proline itself does not influence filament assembly, consistent with the observation that DM3 immune function is independent of catalytic activity. These findings emphasize that structural state—rather than enzymatic output *per se*—underlies DM3’s dual roles in metabolism and immunity. Supporting physiological relevance, mid-sized, curved DM3 filaments are detectable in plant extracts expressing DM3_WT_ and DM3_DT_. Even DM3_WT_, which retains a structurally robust trimer interface *in vitro*, can occasionally adopt filament assemblies *in planta*, indicating that cellular conditions are conducive to structural reconfiguration. The coexistence of multiple processed DM3 forms (5) further suggests that assembly competence may be subject to layered regulatory inputs. Together, these observations point to structural homeostasis of DM3 as a mechanism integrating metabolic adaptation and immune balance. Perturbations that destabilize interface integrities may disrupt this equilibrium, linking supramolecular organization to both stress resilience and immune activation. It would be interesting to directly observe the multivalence status of DM3 *in vivo*. However, capturing this transition *in vivo* remains technically challenging due to the dynamic nature of the hexamer-to-filament conversion. Collectively, our data support a model in which DM3 activity is governed by a reversible transition between an active hexamer and an autoinhibited filament. Through multivalent supramolecular assembly, DM3 integrates environmental responsiveness, metabolic control and immune homeostasis within a unified structural framework.

## Methods

### Protein expression and purification in *E. coli*

DM3 variants were cloned into a modified pET28a vector containing an N-terminal His tag followed by a TEV protease cleavage site. The cloned plasmids were transformed into BL21 (DE3) RILP cells. Cells were grown in LB medium at 37 °C until an OD_600_ of 0.2-0.3, then shifted to 18 °C. Protein expression was induced with isopropyl β-D-1-thiogalactopyranoside (IPTG) at OD_600_ of 0.7-0.8 and incubated overnight at 18 °C. The cultured cells were harvested by centrifugation at 4,000 rpm for 10 min and resuspended in lysis buffer (100 mM NaCl, 50 mM Tris-HCl pH 8.0, 5% (v/v) glycerol). Cells were lysed via sonication and the lysate was centrifuged at 15,000 rpm for 1 h. The supernatant was incubated with Ni-NTA resin (Qiagen) for 1 h at 4 °C. The resin was washed with lysis buffer supplemented with 20 mM imidazole, and proteins were eluted with lysis buffer supplemented with 200 mM imidazole. To remove the N-terminal tag, the eluted proteins were incubated with TEV protease overnight at room temperature. The proteins were further purified by ion-exchange chromatography using a HiTrap Q HP column (Cytiva), followed by size-exclusion chromatography using a Superdex 200 increase 10/300 column (Cytiva).

### Negative-stain electron microscopy

A carbon-coated 1GC400 copper grid (Graticules Optics) was glow-discharged for 20 s at 30 mA using a PELCO easiGlow system (Ted Pella). 3 µL of purified protein (0.03 mg/ml) was applied to the grid and incubated for 1 min. After removing the excess solution by blotting, the grid was washed twice with distilled water. The sample was then stained with 1.5% (w/v) uranyl acetate and air-dried before imaging. Data were collected using a Tecnai F20 transmission electron microscope (FEI) operated at 200 kV at the KAIST Analysis Center for Research Advancement (KARA), Korea.

### Cryo-EM grid preparation and data collection

3 µL of purified DM3_DT_ protein (12 mg/ml in 100 mM NaCl, 50 mM Tris-HCl pH 8.0) was applied to a glow-discharged Quantifoil R2/2 300-mesh copper grid. The grid was vitrified using a Vitrobot Mark Ⅳ (Thermo Fisher Scientific) at 15 °C and 100% humidity, with a blotting time of 3 s and a blot force of −10.

Cryo-EM data were collected using a Titan Krios transmission electron microscope (Thermo Fisher Scientific) operated at 300 kV at the NUS Centre for Bioimaging Sciences (CBIS), Singapore. Movies were recorded with a K3 direct electron detector (Gatan) in electron counting mode at a pixel size of 0.858 Å. Each movie comprised 50 frames with a total exposure dose of 64 e^-^/Å^2^ and was acquired over a defocus range of −2.4 to −0.8 µM. A total of 14,658 movies were collected.

### Filament processing and model validation

Cryo-EM data were processed using CryoSPARC v4.2.1. Beam-induced motion correction and contrast transfer function (CTF) estimation were performed with Patch Motion Correction and Patch CTF Estimation. Of the 14,658 collected micrographs, 12,876 were selected for further processing based on image quality. Filaments were picked using the Filament Tracer tool in a template-free manner. Based on the measured filament diameter (140 Å), the minimum and maximum diameters were set to 70 Å and 180 Å, respectively. Setting the minimum filament diameter to half of the measured filament diameter resulted in optimal filament tracing. Hysteresis thresholds were set to 90 and 95 for low and high thresholds, respectively. A total of 986,130 particles were picked, and 793,154 particles were selected after multiple rounds of 2D classification. An initial reconstruction was performed via asymmetric helical refinement, resulting in a map at 4.17 Å resolution. Helical parameters were determined by a helical symmetry search using information derived from the asymmetric map and 2D classes. Subsequent symmetric helical refinement using these parameters improved the resolution to 3.7 Å, which was further refined to 3.4 Å using non-uniform (NU) refinement. To further improve the map, the 3.4 Å map was used as a reference for template-based picking, yielding 2,220,615 particles. Following the same workflow (2D classification, symmetric helical refinement, and NU refinement), a final reconstruction was obtained at 3.3 Å resolution.

For model building, PDB entry 8JGN was used as the initial model of filamentous DM3. The model was fitted into the 3.3 Å map using ChimeraX, refined by real-space refinement in PHENIX, and manually adjusted in Coot. ChimeraX and PyMOL were used for visualization and figure preparation.

### Prolyl aminopeptidase activity assay

The prolyl aminopeptidase (PAP) activity of DM3 variants were measured using L-proline-7-amino-4-methylcoumarin hydrobromide (Pro-AMC, Merck) as a substrate. 95 µL of 0.5 nM purified DM3 variants were mixed with 5 µL of 20 µM Pro-AMC in a 96-well black plate immediately before measurement. Fluorescence was monitored at an excitation wavelength of 365 nm and an emission wavelength of 440 nm using a BioTek Synergy H1 (Agilent) at room temperature, with readings recorded every 60 s.

### Plant materials

*Nicotiana benthamiana* (*Nb*) plants were grown at 24 °C in a growth room under a 14 h light/10 h dark cycle. Five-week-old plants were used for *Agrobacterium*-mediated transient expression. *Agrobacterium tumefaciens* (strain GV3101) carrying the desired construct was grown overnight at 28 °C in an orbital shaker at 200 rpm in LB medium containing appropriate antibiotics. The bacteria were harvested and resuspended in induction medium (10 mM MES pH 5.6, 10 mM MgCl_2_, and 500 μM acetosyringone) to an OD600 of 1.0. After 3 h of incubation in the dark at room temperature, the *Agrobacterium* culture was mixed with a 1/10 volume of *Agrobacterium* carrying the P19 suppressor of gene silencing. The bacterial mixtures were manually infiltrated into the abaxial side of the leaves using a 1 mL needleless syringe. Infiltrated leaves were kept in the dark for 40 h and then harvested by freezing in liquid nitrogen.

### Plant-derived protein purification from *Nb*

Approximately 20 g of plant extracts expressing FLAG-tagged DM3 variants were harvested 40 h post-infiltration and immediately frozen in liquid nitrogen. The extracts were ground into a fine powder and resuspended in lysis buffer (50 mM NaCl, 50 mM Tris-HCl pH 7.5, 10% (v/v) glycerol, 0.2% NP-40). The resuspended samples were filtered through Miracloth (Merck) to remove coarse cellular debris. The filtrate was clarified by sequential centrifugation at 4 °C, first at 4,000 rpm for 5 min and then 10,000 rpm for 15 min. The supernatant was incubated with anti-FLAG resin (GenScript) for 2 h at 4 °C. After binding, the resin was washed with wash buffer (150 mM NaCl, 50 mM Tris-HCl pH 7.5, 5% (v/v) glycerol, 0.05% NP-40) and proteins were eluted with elution buffer (50 mM NaCl, 50 mM Tris-HCl pH 7.5, 0.4 mg/ml FLAG peptide). The eluate was concentrated using a centrifugal filter device and analyzed by western blot and negative-stain EM.

## Data availability

The cryo-EM map and coordinates are deposited in the EMDB (ID: EMD-69640) and PDB (ID: 24LY), respectively.

## Supporting information

Kimetal_Supple

## Acknowledgements

We thank the KAIST Analysis Center of Research Advancement (KARA) in Korea for negative stain-EM and Glacios cryo-TEM. We thank the NUS Centre for Bioimaging Sciences (CBIS) in Singapore for data collection. We thank the Korea Institute of Science and Technology information (KISTI) and Korea Research Environment Open NETwork (KREONET) for providing computational resources. Use of cryo-EM facilities of CELINE consortium was supported a grant (RS-2024-00440614) by a National Research Foundation of Korea. This work was supported by a grant (RS-2024-00333346, RS-2026-25606952 to J.S.) from National Research Foundation of Korea, the InnoCORE program of the Ministry of Science and ICT (GIST InnoCORE KH0860 to J.S.), Singapore (NRF-CRP22-2019-0001 to E.C.) and Singapore Food Agency (SFS_RND_SUFP_002_04 to E.C.), 2023 Venture Research Program for Graduate & Ph.D. students (VRPGP, No. TRKO202400000904, MSTI to N.K.), and the Brain Korea 21 program (to N.K.).

## Author contributions

N.K., E.C., and J.S. conceived the project. N.K. performed the biochemical and structural experiments, including negative-stain EM, cryo-EM data processing, model building and analysis. W.-L.W. and Y.-Y.T. performed agroinfiltration and provided plant extracts used in this study. N.K., E.C. and J.S. wrote the manuscript. All authors examined the data and revised the manuscript. E.C. and J.S. supervised the project and acquired funding.

---

**Table 1.**
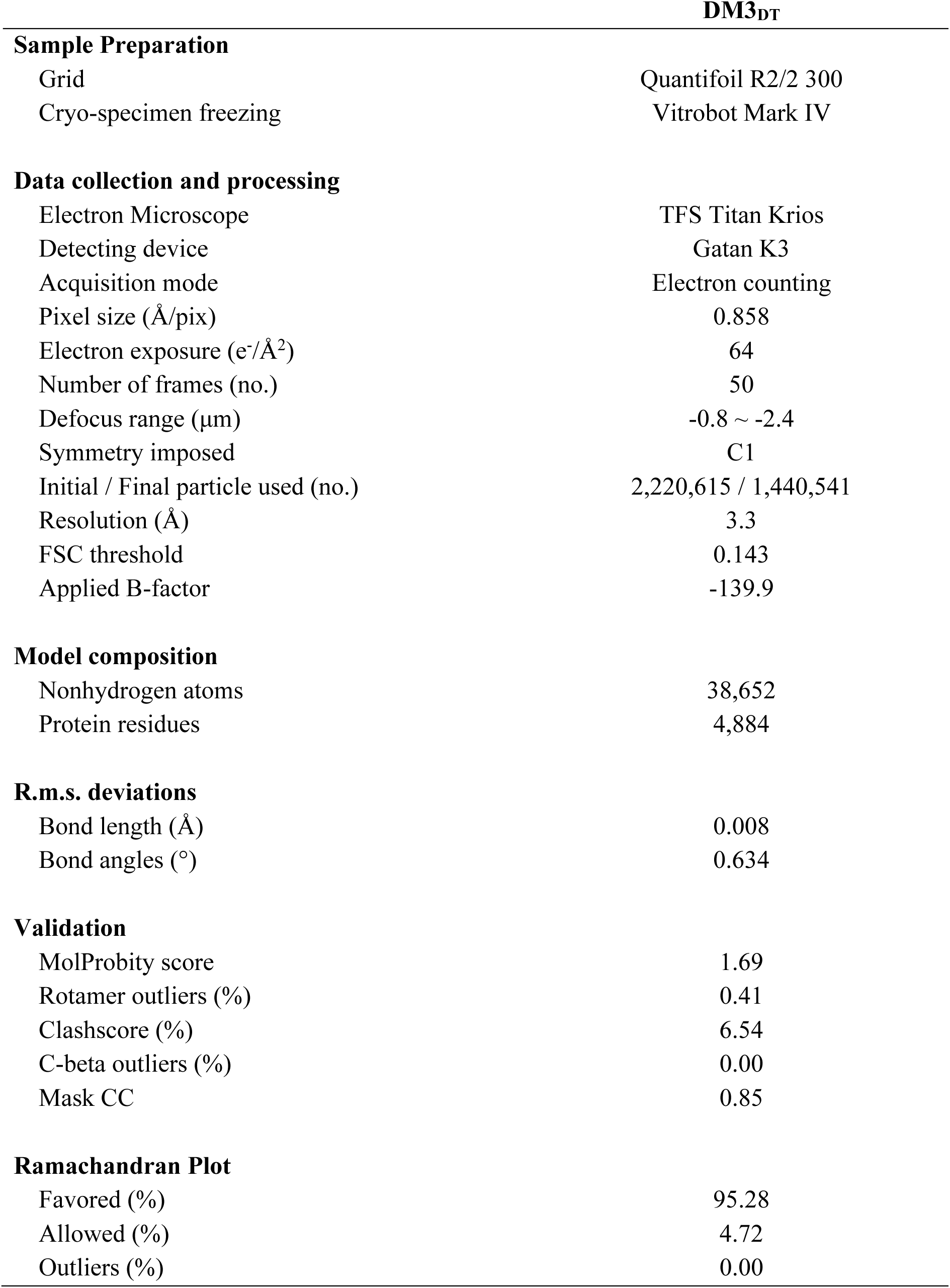
Cryo-EM sample preparation, data collection and model validation.

