## Supplementary material for "Dynamic Filament Assembly Regulates the Prolyl Aminopeptidase Activity of Plant Immune Protein DM3": Kimetal_Supple

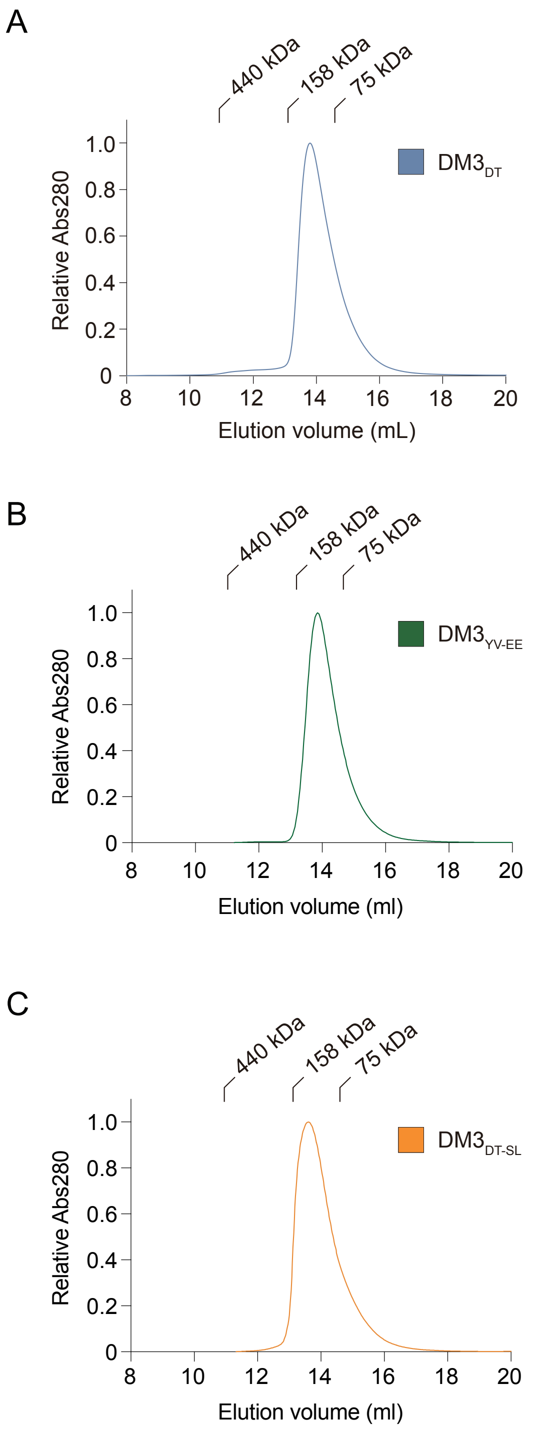


**Figure S1.** Size-exclusion chromatograms of DM3 variants. *(A, B, and C)* Size-exclusion chromatogram of $DM3_{DT}$ *(A)*, $DM3_{YV\text{-}EE}$ *(B)*, and $DM3_{DT\text{-}SL}$ *(C)*. Standard molecular weight markers are indicated at the above of the chromatograms.

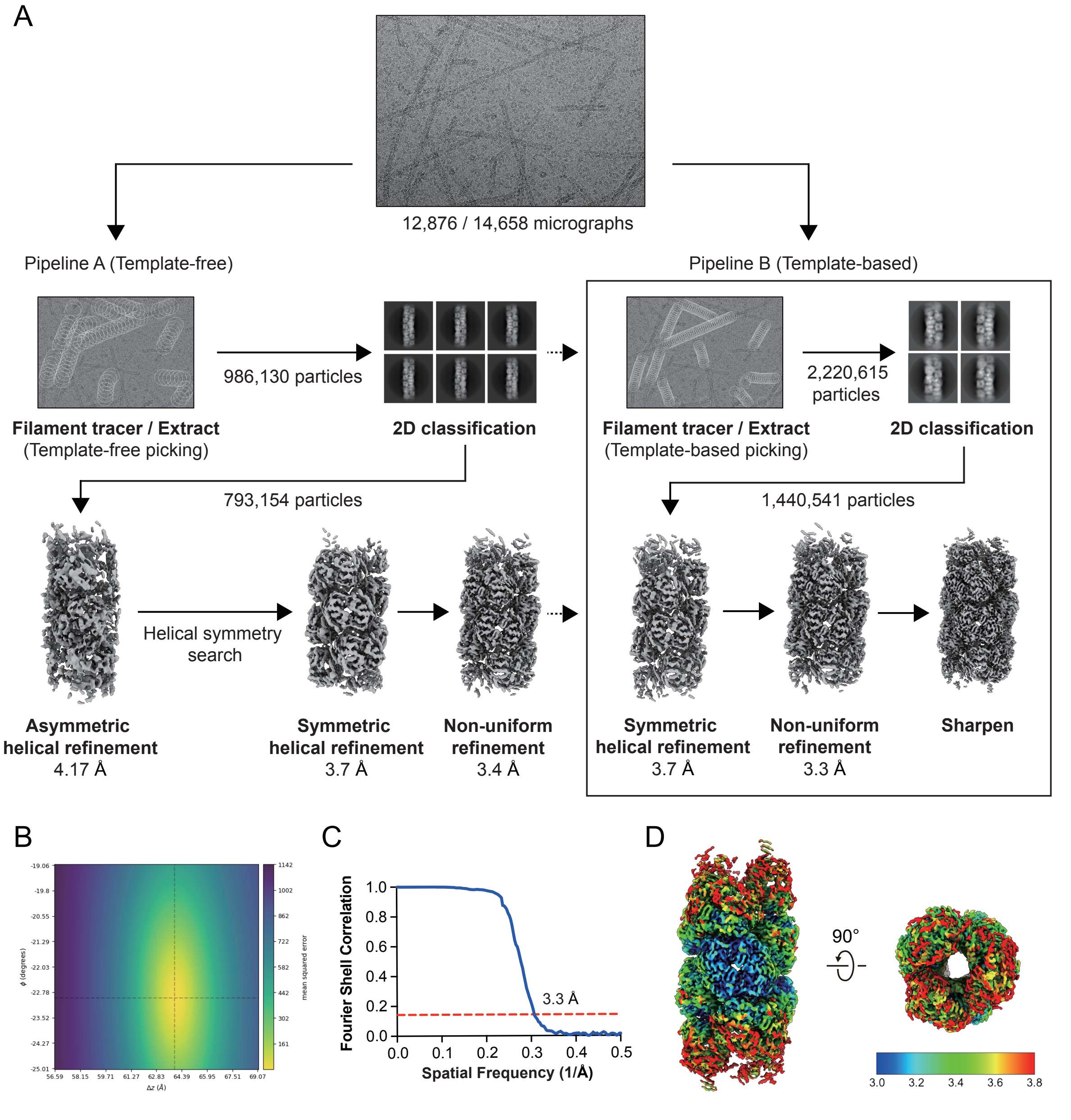


**Figure S2.** Cryo-EM data processing of $DM3_{DT}$ filament structure. *(A)* A flowchart of the cryo-EM data processing. Pipeline A performed template-free picking for the initial reconstruction, which then served as a template for Pipeline B to perform template-based picking. Dash arrows indicate that the 2D classification and non-uniform refinement volume from pipeline A serves as templates for filament tracing and symmetric helical refinement in pipeline B, respectively. *(B)* CryoSPARC symmetry search identified a peak at a rise of 63.96 Å and a twist of -22.8°. These parameters were used for helical refinement, yielding a final converged rise of 63.9 Å and a twist of -22.5°. *(C)* The FSC curve shows a resolution of 3.3 Å for the $DM3_{DT}$ filament at the 0.143 criteria. *(D)* Local resolution of the cryo-EM map of $DM3_{DT}$ filament is estimated by CryoSPARC and colored by ChimeraX.

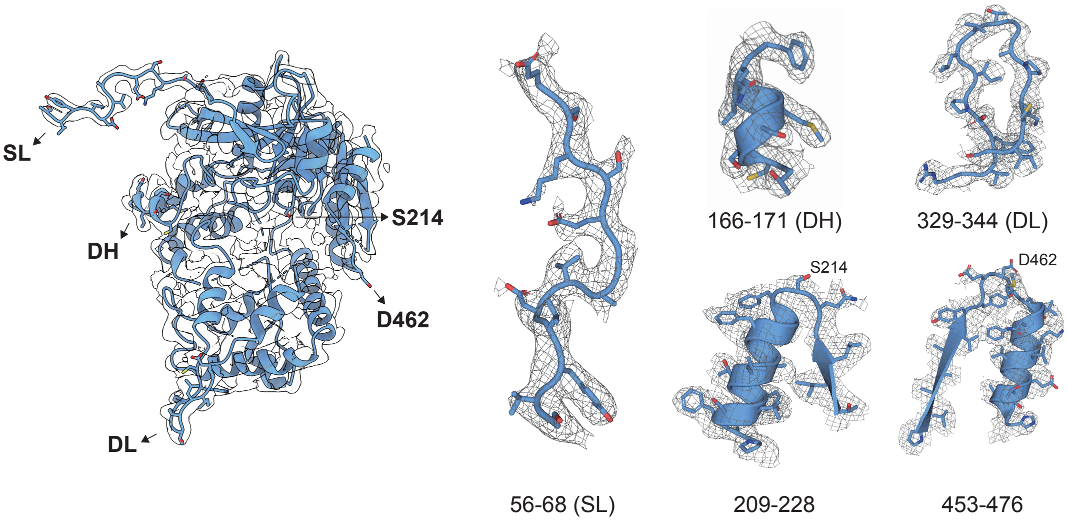


**Figure S3.** Representative cryo-EM density with fitted atomic model of $DM3_{DT}$. The cryo-EM structure of $DM3_{DT}$ is fitted into the cryo-EM map using PyMOL. The stacking helix (SL), Dimerizing Helix (DH), Dimerizing Loop (DL), and regions encompassing the catalytic triad components S214 (residues 209-228) and D462 (residues 453-476) are shown.

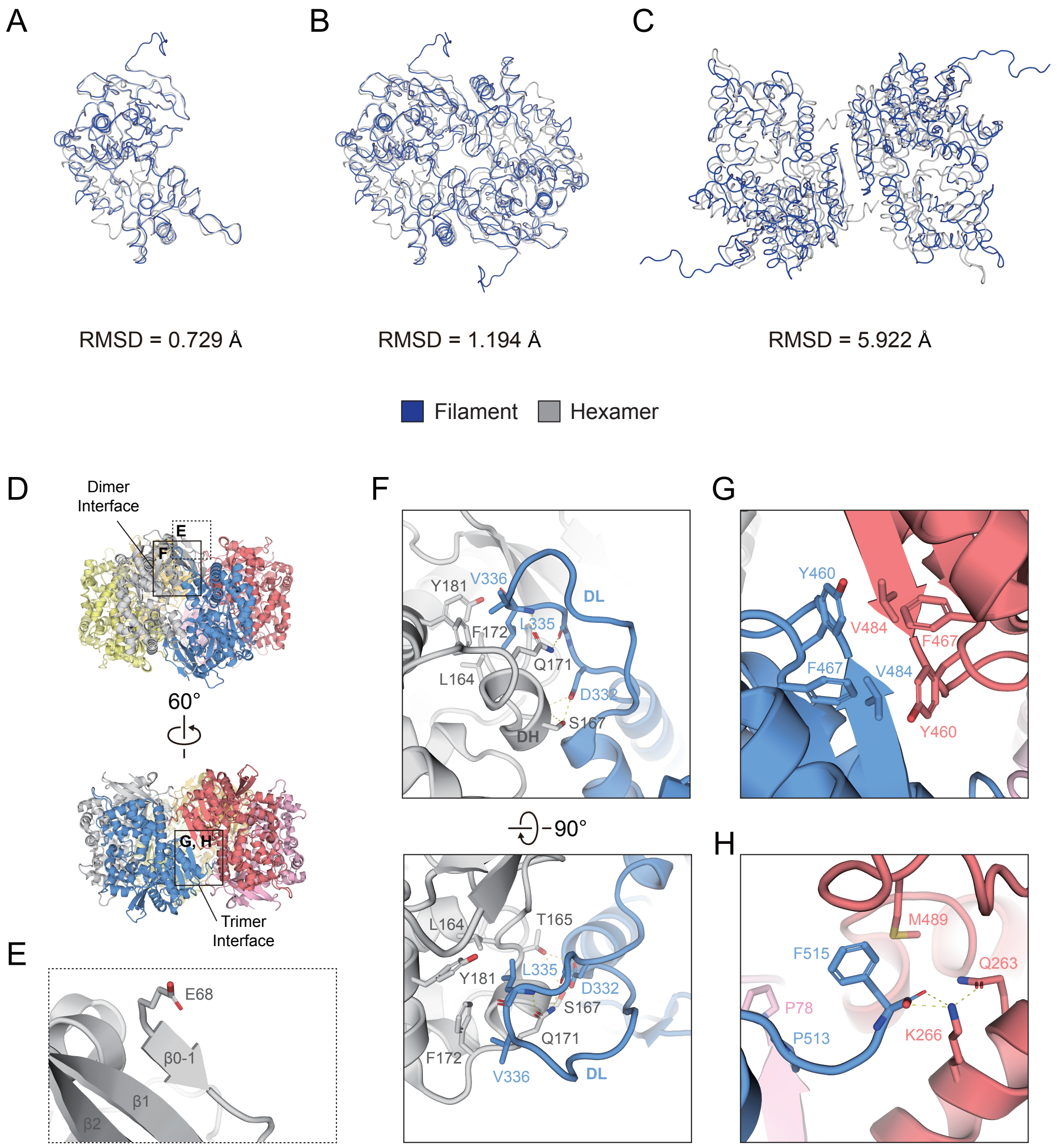


**Figure S4.** Structural comparison between the planer and filamentous DM3. *(A)* Monomers from the filament (blue) and hexamer (gray) superimpose with an RMSD of 0.729 Å. *(B)* Two adjacent DM3 subunits at the dimer interface from the filament (blue) and hexamer (gray) superimpose with an RMSD of 1.194 Å. *(C)* Two adjacent DM3 subunits at the trimer interface from the filament (blue) and hexamer (gray) superimpose with an RMSD of 5.922 Å. *(D)* Dimer and trimer interface are specified on the cryo-EM structure of hexameric DM3. Each interface is indicated by a box. The region encompassing residue E68 is indicated by a dashed box. *(E)* Enlarged view of the region encompassing residue E68. E68 is exposed on the outer surface. *(F)* Detailed view of dimer interface formed by DH and DL. *(G)* Detailed view of the trimer interface highlighting hydrophobic interactions. *(H)* Detailed view of the trimer interface highlighting the C-terminal insertion.

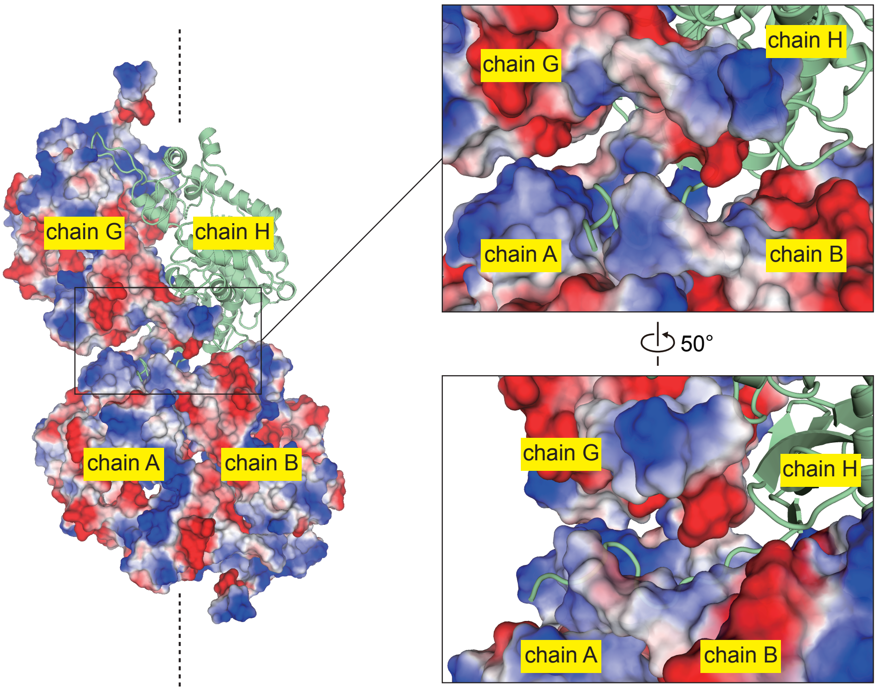


**Figure S5.** Insertion of the stacking loop into the DH-DL pocket. Surface representation of the filament colored by electrostatic potential (-50 to +50 mV). The filament axis is indicated by a dotted line. For visualization, chain H is shown as a cartoon, highlighting the insertion of the stacking loop into the DH-DL pocket.

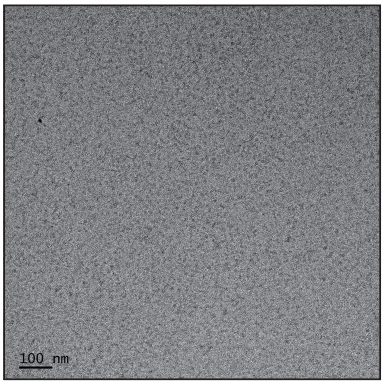


0 mM NaCl

**Figure S6.** Representative negative-stain EM image of $DM3_{DT}$ at 0 mM NaCl, showing no detectable filament formation.
